# Diversity and ecological potentials of viral assemblages from the seamount sediments of the Northwest Pacific Ocean

**DOI:** 10.1101/2024.04.28.591573

**Authors:** Ying Chen, Chen Gao, Qian Liu, Yantao Liang, Mingyan Lai, Fuyue Ge, Hao Yu, Hongbing Shao, Andrew McMinn, Min Wang

## Abstract

Viruses are the most abundant life forms in the sea, influencing the community structure and metabolism of host cells as well as biogeochemical cycles. However, the diversity and ecological roles of viruses within seamount ecosystems, natural microbiota havens characterized by high biodiversity, remain unknown. Here, the first seamount viral genome (SMVG) dataset, based on a metagenomic analysis of twelve seamount sediment samples collected from the seamount regions of the Northwest Pacific Ocean, is established. A total of 78,069 viral operational taxonomic units (vOTUs) were found, spanning 18 viral classes and 63 viral families. The detection of sixteen viral auxiliary metabolic genes (vAMGs) suggests that viruses may participate in both the complex metabolic dynamics associated with sediment microbial communities and also biogeochemical cycles, including carbon, sulfur, metal, heme, and cobalamin cycling. vAMGs involved in the metabolism of heme, cobalamin and metals, in particular, are more often detected in seamount sediments than in trenches, cool seeps, and hydrothermal vents. This investigation of the viral communities in these seamount sediments provides new insights into the high diversity and ecological potential of the viruses and establishes a foundation for the future study of benthic viruses from deep-sea seamounts.

## Introduction

Many deep-sea habitats, such as sea mountains, cold seeps and hydrothermal vents, have particularly high biodiversity. Within these habitats, viruses might have evolved numerous strategies to ensure their successful reproduction. This includes their ability to reprogram the hosts’ metabolism via the viral auxiliary metabolic genes (vAMGs), horizontal gene transfer, etc. [1–4]. Importantly, as critical mortality agents of both prokaryotes and eukaryotes, they can also affect biogeochemical cycles and ecosystem dynamics [5, 6]. Benefiting from rapid advances in metagenomics, viral metagenomic data presents an opportunity for large-scale identification of unclassified viruses from the deep ocean, from areas such as trenches, cold seeps, and hydrothermal vents [7–9].

Seamounts are isolated topographic features with summit heights at least 100 m above the seafloor but, not reaching the sea surface [10–12] Seamounts are generally volcanic in origin and worldwide are recognized as important offshore deep ecosystems [13]. These structures are frequent on the oceanic crust and transitional crust and interact with ocean currents, triggering complex oceanographic processes, such as the “Taylor column”, turbulent mixing and benthic boundary effects. [14, 15]. Although there are around 33,452 identified seamounts and 138,412 knolls in the global ocean, only 1.5% of seamounts and 0.7% of knolls are located within marine protected areas listed on the World Database of Protected Areas (WDPA) [16]. Currently, less than 300 seamounts have been explored [17], primarily in the Northwest Pacific, leaving a dearth of information from other areas.

Recent investigations of seamount ecology have shown that their geographical features and hydrological conditions allow for the existence of abundant prokaryotes and their predators, including pelagic marine predators and viruses, which then release labile organic matter into the surrounding seawater and sediments [8, 14]. Biological and ecological studies over many years have shown that prokaryotic abundance in sediments surrounding seamounts is typically 2-3 times higher than in non-seamount sediments [14]. These bacteria and archaea, which are chemolithoautotrophic, use sulfur, nitrate, and heavy metals as energy sources [8, 18–20] and may even have developed morphological adaptions to the seamount environment [21]. Notably, viruses have been demonstrated to manipulate the mortality of microorganisms, releasing labile organic matter, and expressing vAMGs that can rewire host metabolism [6, 8, 22]. However, the community composition, diversity and ecological roles of viral assemblages inhabiting seamount sediments are still unknown.

In this study, viral communities from Northwest Pacific Ocean seamount sediments, together with their metabolic potential, were investigated. The first seamount viral genome (SMVG) dataset was established and the viral community structure in the sediment surface viral communities was examined using metagenomic technology and a comprehensive analysis of the genomic diversity, distribution, lifestyle, and ecological function.

## Materials and Methods

### Sample collection

Twelve surface sediment samples were collected from Northwest Pacific Ocean seamounts, within a geographic range of 148.43942°E to 156.522634°E and 12.429233°N to 23.23131°N, over three years (Supplemental Table. S1). The sediment samples were collected from the top 1 cm of the surface layer using a box corer (DY61_MC0201 using multi-corer). The collected samples were immediately frozen at -80°C until further processing.

### DNA extraction and sequencing

The total DNA from sediment samples (0.5 g) from cruise DY48 was extracted by Novogene (Nanjing, China), DNA from cruises DY56 and DY61 was extracted by Guangdong Magigene Biotechnology Co., Ltd. (Guangzhou, China), using ALFA-SEO Magnetic Soil DNA Kit according to the manufacturer’s instructions, respectively. DNA integrity and purity were monitored on 1% agarose gels. DNA concentration and purity were measured using Qubit 3.0 (Thermo Fisher Scientific, Waltham, USA) and Nanodrop One (Thermo Fisher Scientific, Waltham, USA) at the same time. Subsequently, the total DNA of each sample (100–200 ng DNA as a template) was amplified with QIAGEN REPLI-g Mini Kit (phi-29 DNA polymerase), which used the whole genome multiple displacement amplification (MDA) method (details see Supplemental Information) [23, 24]. Library preparation and high-throughput sequencing of each sample were sequenced on the Illumina platform and 150 bp paired-end reads at the Novogene (Nanjing, China) or using NEB Next® Ultra™ DNA Library Prep Kit for Illumina® (New England Biolabs, MA, USA) following the manufacturer’s recommendations and index codes were added at the Guangdong Magigene Biotechnology Co., Ltd. (Guangzhou, China).

### Quality control and assembly and identification

Raw reads underwent quality control by removing adapters using Fastp (-n 0) [25]. Subsequently, high-quality paired-end reads were filtered by Perl scripts, ensuring the following criteria: (1) absence of N; (2) no more than 20% of bases with a quality score below 20; (3) no more than 30% of bases with a quality score below 30 [26, 27]. Quality-controlled reads from each sample were assembled using MetaHIT v1.2.9, with the removal of contigs ≤ 1,500 bp before viral identification [28, 29], then metagenomic assemblies were evaluated by metaQUAST (default mode). Metagenomic contigs (> 1,500 bp) from each site were subjected to viral identification tools: VirSorter v1.0.6, VirFinder. The following criteria were applied for viral contig identification: (1) contigs sorted by VirFinder with a score ≥ 0.9 and p < 0.05; (2) contigs sorted by VirSorter in categories 1 and 2; (3) contigs sorted by VirFinder with a score ≥ 0.7 and p < 0.05 and VirSorter categories 1-6. CheckV v0.7.0. was used to remove non-viral fragments and mitigate the effects of microbial contamination. .

### Dereplication, relative abundances, and taxonomic profiling

Viral contigs from each sample were dereplicated using CD-HIT v4.8.1 (parameters: - c 0.95 -G 0 -M 0 -aS 0.8 -T 8 -n 9), meaning that viral contigs met the condition that ≥ 95% nucleotide identity across ≥ 80% of shorter viral contigs were considered to be the same viral population and were removed [30]. As a result, 78,069 viral operational taxonomic units (vOTUs) were retained. TPM (Transcripts per million) values were used to represent the relative abundance of viruses. Quality-controlled contigs from each sample were analyzed using CoverM v0.6.1 to count coverage profiles across samples (parameters: -p bwa-mem --min-read-percent-identity 0.95 --min-read-aligned-percent 0.75 -m tpm -t 20) [31, 32]. VITAP was used to place the 78,069 vOTUs in taxonomic profiling, which was based on the fitting of sequence similarity and the classification extension based on the bipartite graph, and the lifestyle of viruses was detected by VIBRANT v1.2.1 with the default parameters (except - virome) [33].

### Phylogenetic analysis

The viral terminase large subunit (Terminase_1, Terminase_3, Terminase_6N, Terminase_Gpa) and the NCLDV-derived family B DNA polymerase (*PolB*) protein were used as the conserved marker genes to construct phylogenetic trees of *Caudoviricetes* and Nucleocytoplasmic large DNA viruses (NCLDVs) [34]. For the phylogenetic trees of viral terminase, a total of 990 viral terminase large subunits (TerL) were identified from the samples by PfamScan. Then a total of 792 reference sequences were downloaded from the NCBI virus database (NCBI Virus (nih.gov)) and a dereplication was performed (Supplemental Table S3). The NCLDV *polB* proteins were searched against a hidden Markov model of NCLDV *PolB* sequences using hmmsearch (-E 1e-10). Reference sequences of the *polB* protein were obtained from a previous study [35]. Sequences with less than 300 amino acids were removed.

The 36 dereplicated sequences, together with 500 reference sequences, were aligned using MAFFT v7.490 (auto mode), then trimmed using TrimAL (v1.4. rev15) to remove position with gaps over 90% [36, 37]. The maximum-likelihood phylogenetic trees were constructed using IQ-TREE v2.1.4 (--safe -m MFP -B 1000) [38] and visualized using iTOL (version 6) [39].

### Comparison with viruses from other habitats

To understand the similarities and differences between viruses from seamounts sediments and other environments (including oceanic, deep subsurface, non-marine saline and alkaline environments, and extreme habitats), 2,890 viral contigs with lengths over 10 kb were selected from SMVG, subsequently, 20,195 viral reference sequences were used to construct the dataset, including 4,509 high-quality viral genomes from oceanic habitats (IMG/VR v4), 4,060 high-quality viral genomes from the deep subsurface (IMG/VR v4), 3,379 high-quality viral genomes from non-marine saline and alkaline environments (IMG/VR v4) and 8,247 viral sequences from other extreme habitats. Specifically, the ‘extreme habitats’ consisted of 1,626 sequences from hydrothermal vents, 2,885 sequences from cool seeps [40], and 3,736 sequences from Challenger Deep [8]). For each viral contig, ORFs were predicted using metaProdigal v2.6.3 [41], then all-verses-all DIAMOND v0.9.29.130 BLASTP was used to compare the protein with parameters: e-value ≤ 1 ⅹ 10^-5^, query coverage ≥ 50%, identity ≥ 25% [29, 42]. All sequences were analyzed using vConTACT 2 with the following parameters: –db ‘None’ –pcs-mode MCL –vcs-mode ClusterONE. The network is illustrated using Cytoscape v3.9.1 and the number of VCs was plotted using Evenn [43, 44].

### Host binning and taxonomic profiling

Quality-controlled metagenomic sequences were binned (parameters: --metabat2 -- maxbin2 --concoct) and refined (parameters: -c 70 -x 5) using metaWRAP v1.3.2 [40, 45]. A total of 155 metagenome-assembled genomes (MAGs) were initially produced, and these were then de-replicated into 71 using dRep v3.2.0 (parameters: -sa 0.95 -nc 0.30 -p 24 -comp 70 -con 10) [46]. Taxonomic annotation of the MAGs was carried out using GTDB-Tk v1.0.2 [47], and relative abundance was calculated by TPM using CoverM v0.6.1 (parameters: -p bwa-mem --min-read-percent-identity 0.95 --min-read-aligned-percent 0.75 -m tpm -t 20).

### Virus-host prediction

Four different *in silico* methods were employed to predict virus-host interactions,[40]. (1) Nucleotide sequence homology; vOTUs were compared with sequences of MAGs using BLASTn with parameters: query coverage ≥ 75%, identity ≥ 70%, bit score ≥ 50, and e-value ≤ 1e-3; (2) Oligonucleotide frequency (ONF); oligonucleotide frequency and distance between vOTUs and MAG sequences were calculated using VirHostMatcher v1.0 with default parameters. Predictions with d2* values ≤ 0.2 were considered eligible for assigning virus-host linkages [48]. (3) Transfer RNA (tRNA) match; screening of tRNA genes from viral contigs and MAGs was performed with ARAGORN v1.2.38 with the parameters “t” [49]. A match was identified when there was ≥ 90% length identity in ≥ 90% of the sequences by BLASTn [50]. (4) CRISPR spacer match; CRISPR arrays were assembled from clean reads of metagenomes using class v1.0.1 with default parameters [51]. The retained spacers from reads were then compared with vOTUs using BLASTn (e-value ≤ 10^-5^, mismatch ≤ 1) [52]. Replicate comparisons were performed using BLASTn with the same parameters as for prokaryotic MAGs, for each matched CRISPR spacer, [40].

### Functional annotation and abundance of ORFs and putative vAMGs

The ORFs from viral contigs were identified using metaProdigal with default parameters. A comprehensive annotation of viral ORFs was performed based on four databases, CAZy, eggNOG, KEGG and pVOG databases [53]. Firstly, ORFs were compared with the Carbohydrate-active enzymes (CAZy) database using DIAMOND (parameters: (e value 1ⅹ10^-5^; > 60% identity)), to identify the carbohydrate-metabolism-related gene [54]. Furthermore, all ORFs were annotated against the eggNOG v5.0 database to classify COG functional classifications by eggnog-mapper with default parameters [55]. Additionally, the ORFs were submitted to GhostKoala to be annotated by the KEGG database (https://www.kegg.jp/ghostkoala/) [56]. All the ORFs were compared with the Prokaryotic Virus orthologous groups (pVOGs) database[53], then ORFs with >40% viral regions and >15% viral hallmarks and viral-like genes were manually selected. Subsequently, the vAMGs were manually identified based on the criteria that both the start and end genes were annotated as viral proteins by the pVOG database. The relative abundance of putative vAMGs in each site was calculated by CoverM v0.6.1 (parameters: -p bwa-mem --min-read-percent-identity 0.95 --min-read-aligned-percent 0.75 -m tpm -t 20). Heatmaps were generated and visualized using TBtool v1.108 [57].

For the phylogenic analysis of cobaltochelatase (*cobS* and *cobT*), reference protein sequences were retrieved from IMG/VR v4 [58] and the UniProt databases. *cobS* and *cobT* reference sequences were retrieved from the IMG/VR v4 database by BLASTp with parameters: e-value ≥ 1e-10, 244 and 475 high-quality sequences were retrieved, respectively. Subsequently, reference sequences from the IMG/VR database were de-replicated using CD-HIT v4.8.1 (parameters: -c 1 -G 0 -M 0 -aS 0.8 -T 8 -n 9), 119 and 138 viral reference protein sequences were retained from IMG/VR v4 database. Microorganism reference sequences were downloaded from the UniProtKB database, resulting in a total of 247 and 4,766 reference sequences, respectively. Reference sequences from the UniProtKB database were de-replicated using CD-HIT v4.8.1, then 204 and 193 microorganism reference sequences were retained from the UniProtKB database. The construction of a phylogenetic tree is described in the section “Phylogenetic analysis”.

### Comparison to specific vAMGs from other habitats

To compare SMVG viral heme, cobalamin and metal vAMGs with those from other habitats, three ecological environmental metagenomic data sets were used, including Treachs [8], hydrothermal vents [59], cool seeps [40] (samples detail see Table S9), and data from this study comprising 79 samples in total. Contigs associated with heme, cobalamin and metal functional annotations from the KEGG database were singled out, and then passed to CoverM v0.6.1 to count coverage profiles across metagenomic data sets (parameters: -p bwa-mem --min-read-percent-identity 0.95 --min-read-aligned-percent 0.75 -m tpm -t 20).

### Statistical Analysis

All statistical analyses were performed using R version 4.2.3. vegan v2.6.4 was used to calculate the alpha diversity based on the relative abundance of viral contigs [60]. For comparison of different environments (Table S9), i.e. seamount sediments, cold seeps, hydrothermal vents and hadal trenches, all viral contigs with lengths over 10 kb were selected to calculate the Shannon and Simpson diversities.

### Results and Discussion

Most previous studies on seamounts and deep sea sediments have focused on components of the microbial ecosystems [21, 61, 62]. These studies are complemented here by studying the viral community of seamount sediments. Here, the distinct viral biogeographic distribution patterns, the relationship between viruses and their hosts, endemism, functional genes, and biosynthetic potentials of viral assemblages inhabiting the seamount sediments are explored for the first time.

### Overview of viral community composition in the seamount sediments

This first seamount viral genome (SMVG) dataset was assembled from 12 DNA virome libraries with a total of 225.4 GB of sequence data across 12 seamount sediment samples. Database assembly yielded 29.97 GB of sequence data (Fig. 1A). From this dataset, 81,418 viral contigs with a size ≥ 1.5 kb were retrieved and classified into 78,069 viral Operational Taxonomic Units (vOTUs). Of these, 444 and 244 viral contigs were identified as complete or high-quality genome sequences, respectively (Supplemental Table S2). The most abundant vOTUs differed between the three cruises (Fig. S1) and the five viromes of DY56 have the highest number (n=55). Based on the VIBRANT results, 35,510 of the 78,069 vOTUs (∼45.49%) were predicted to be lytic viruses, and 716 (∼0.91%) were predicted to be temperate viruses (Fig. S2).

**Fig. 1.**
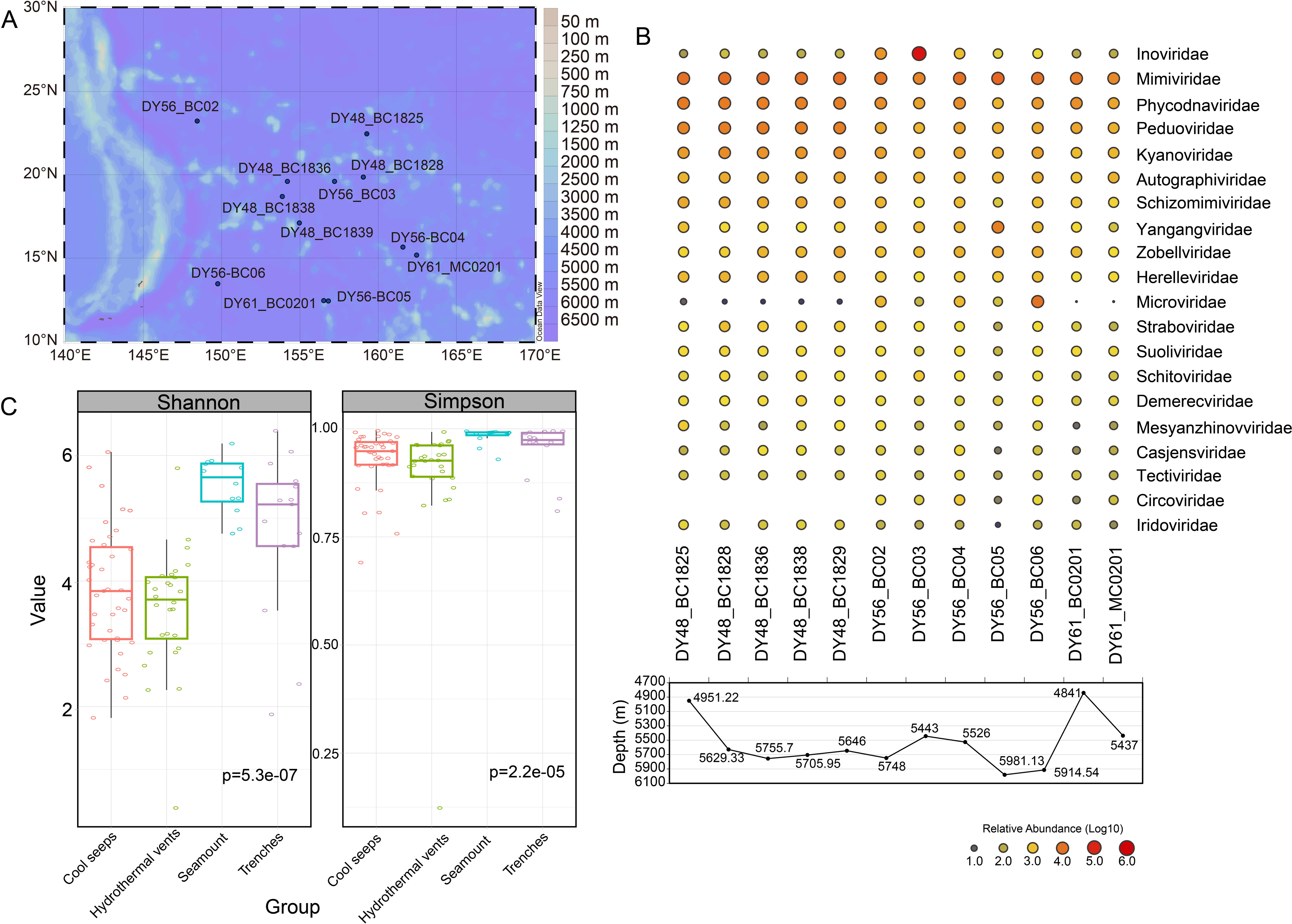
Overview of the seamount sediments viruses distribution and identified in this study. **(A)** Map of sampling of the seamount sediment in the Northwest Pacific Ocean in this study (different latitudes were tinted by different colors). **(B)** Relative abundance of viral contigs at the top 20 family level (upper half) and the water depth of sampling stations. (lower half). **(C)** Shannon and Simpson indices of the viral community from different environments. Relative sequence abundance was calculated using CoverM v0.6.1.

A total of 17,794 vOTUs (∼22..79%) were taxonomically assigned to 63 families and 18 to class level. *Inoviridae* (∼7.71%), *Mimiviridae* (∼5.42%), and *Phycodnaviridae* (∼2.65%) had the highest relative abundance (Fig. 1B). Of the top 20 most abundant families, 12 belonged to class *Caudoviricetes*, with no noteworthy distinction between sites. (Fig. 1B). By contrast, *Microviridae*, *Circoviridae*, *Smacoviridae* and *Vilyaviridae*, which are icosahedral viruses with a circular single-stranded DNA genome, were mainly detected in sediments of the DY56 cruise (Fig. S3).

A comparison of Shannon diversity and Simpson diversity indicies of the viral community revealed different diversities between the four environments, i.e. cold seeps (n= 2885), hydrothermal vents (n=935), seamounts (n=2883) and hadal trenches (n= 3736), with seamount sediment having a higher viral diversity than the other three environments (Fig. 1C).

### Phylogenetic diversity of *Caudoviricetes* and NCLDV includes novel and ancient viruses

The phylogenetic tree predominantly grouped the seamount sediment viruses as unclassified groups within the *Caudoviricetes* class (Fig. S4). Different types of TerL domains were located on significantly different branches and the sequences in the samples were widely distributed among the various types of TerL domains.

Interestingly, the terminase sequences formed a distinct and divergent group with long branches (Fig. S4). Furthermore, the presence of certain terminase-based viral clades solely in the samples investigated here and located near the root of the phylogenetic tree, implies that they could potentially represent endemic and ancient *Caudoviricetes* taxa in the seamount sediments.

Furthermore, nucleocytoplasmic large DNA viruses (NCLDVs) showed high relative abundance in the 12 samples, thus, viral proteins were matched to the *polB* marker gene using a hmmsearch search, which detected 36 *polB* sequences in the samples (Supplemental Table S4). Unlike the phylogenetic tree of the TerL domain, a large proportion of the *polB* sequences formed an independent branch, except for reference sequences, indicating that the majority of NCLDVs in the seamount sediments might have a similar phylogenetic status but were different from known NCLDVs (Fig. S5). As reference sequences were collected from shallow or intermediate water depths (1-1000 m) [63] and 30 *polB* seamount sediment sequences have a close phylogenetic position with reference sequences, it was hypothesized that the deep-sea viruses could be linked to the surface ocean [64, 65]. However, this hypothesis remains unconfirmed.

### Comparison of viral endemism between seamount sediments and other ecosystems

To investigate the relationship between seamount vOTUs and publicly available virus sequences from a wide number diversity of ecosystems, a gene-sharing network was constructed using vConTACT 2 (Fig. 2A). Seamounts from this study, deep subsurface, oceanic, non-marine saline and alkaline and extreme ecosystem were grouped into 3,345 viral clusters (VCs). Surprisingly, only 15 VCs were shared amongst all ecosystems (Fig. 2B). Different virus community structures were observed among the 11,982 vOTUs identified from the seamount sediments. 580 VCs were formed, of which 233 were exclusively found in this ecosystem and not detected elsewhere. This implies that ∼40.17% of total vOTUs in seamount sediments were absent from the other four habitats included in the IMG/VR v4 dataset. In addition, the presence of novel and endemic viruses was evident in the seamount sediments, with each station containing distinctive vOTUs, with DY56_BC05 having the highest number of unique vOTUs (Fig. 2C). Deep-sea seamounts are generally covered with ferromanganese (Fe–Mn) crusts [66–68]. These Fe-Mn crusts and nodules support microbial communities and play a role in the cycling of Fe and Mn and the growth of Fe-Mn crusts [61]. Therefore, the presence of cobalt crusts, identified on cruise DY56, may have contributed to the endemism of viruses in the seamount sediments.

**Fig. 2.**
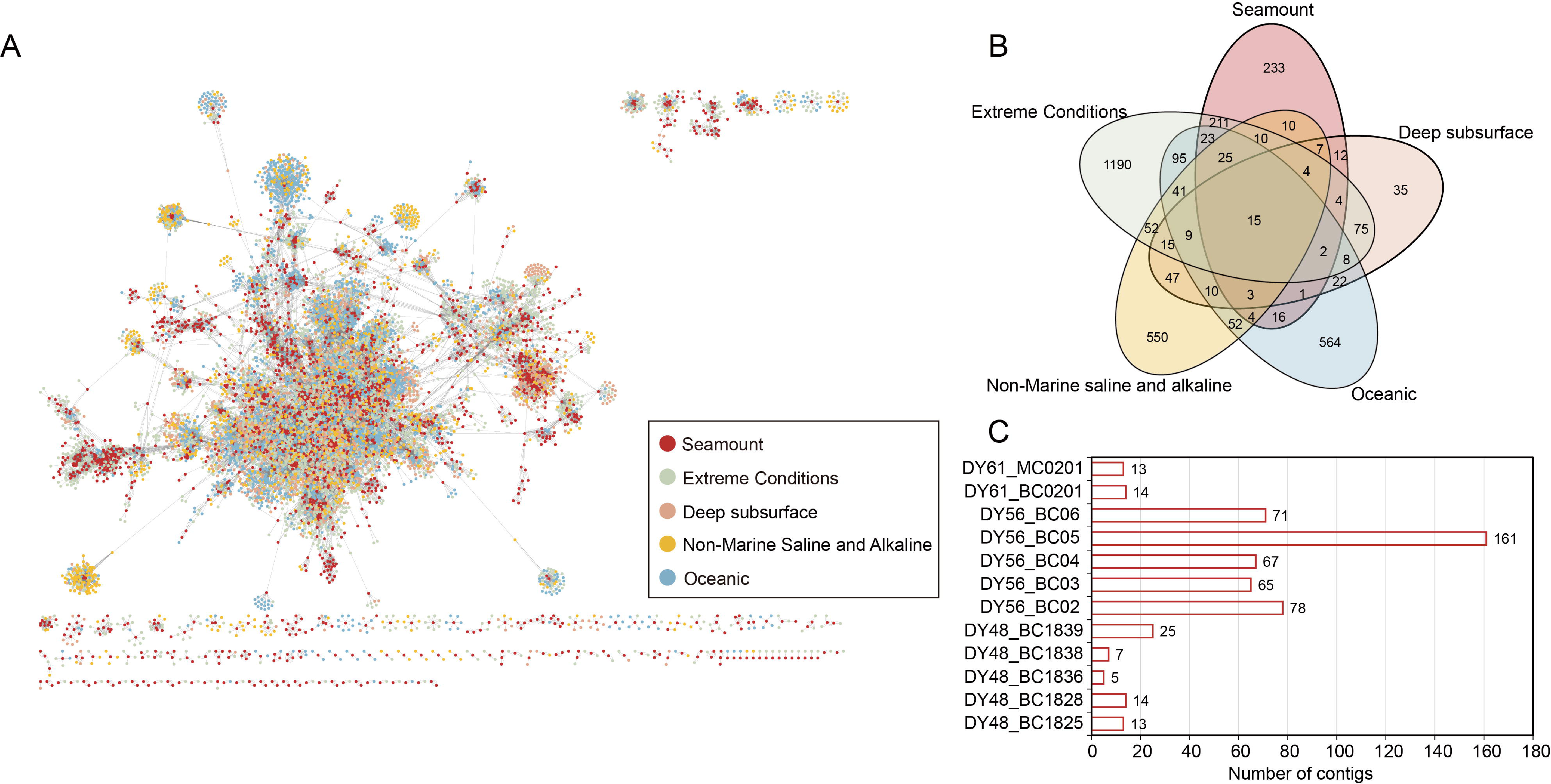
Comparison analysis between seamount sediments vOTUs and other environmental data sets. **(A)** The gene-sharing network shows the comparison among five environmental viral contigs, five habitats including seamount sediments, extreme conditions (i.e. challenger deep, cold seep, and hydrothermal vents), deep subsurface, non-marine saline and alkaline, and oceanic. **(B)** Venn diagram of shared viral clusters (VCs) among five environments. **(C)** Bar charts of specific viral contigs among different stations based on **(B)**.

### Diversity and virus-host linkages in seamount sediments

To explore the association between viruses and their potential hosts, an analysis involving sequence similarity comparison, oligonucleotide frequencies, tRNA sequences and CRISPR spacers was performed [69]. However, most potential vOTU hosts could not be classified, and only 843 vOTUs of those present in the seamount sediments (1.14% of the total vOTUs) were linked to twenty-six potential hosts (Table S6).

Overall, only 30 vOTUs exhibited a broad host range, with the majority (813) displaying a narrow host range (Fig. 3A), consistent with previous observations [40, 70, 71]. Even though 98.86% of the contigs identified in this study could not be linked to a host, a significant positive correlation between viruses and hosts, based on the matching of the relative abundance of host-linked viruses (*P* < 0.001), was observed, supporting the previously predicted result that the composition of predicted microbial hosts aligns well with the composition of their viruses (Fig. 3B). Additionally, when examining the proportion of host-linked viruses with lysogenic and lytic lifestyles using VIRBRANT, most of the viruses with potential hosts (171 vOTUs) were found to have a lytic lifestyle (Fig. 3C).

**Fig. 3.**
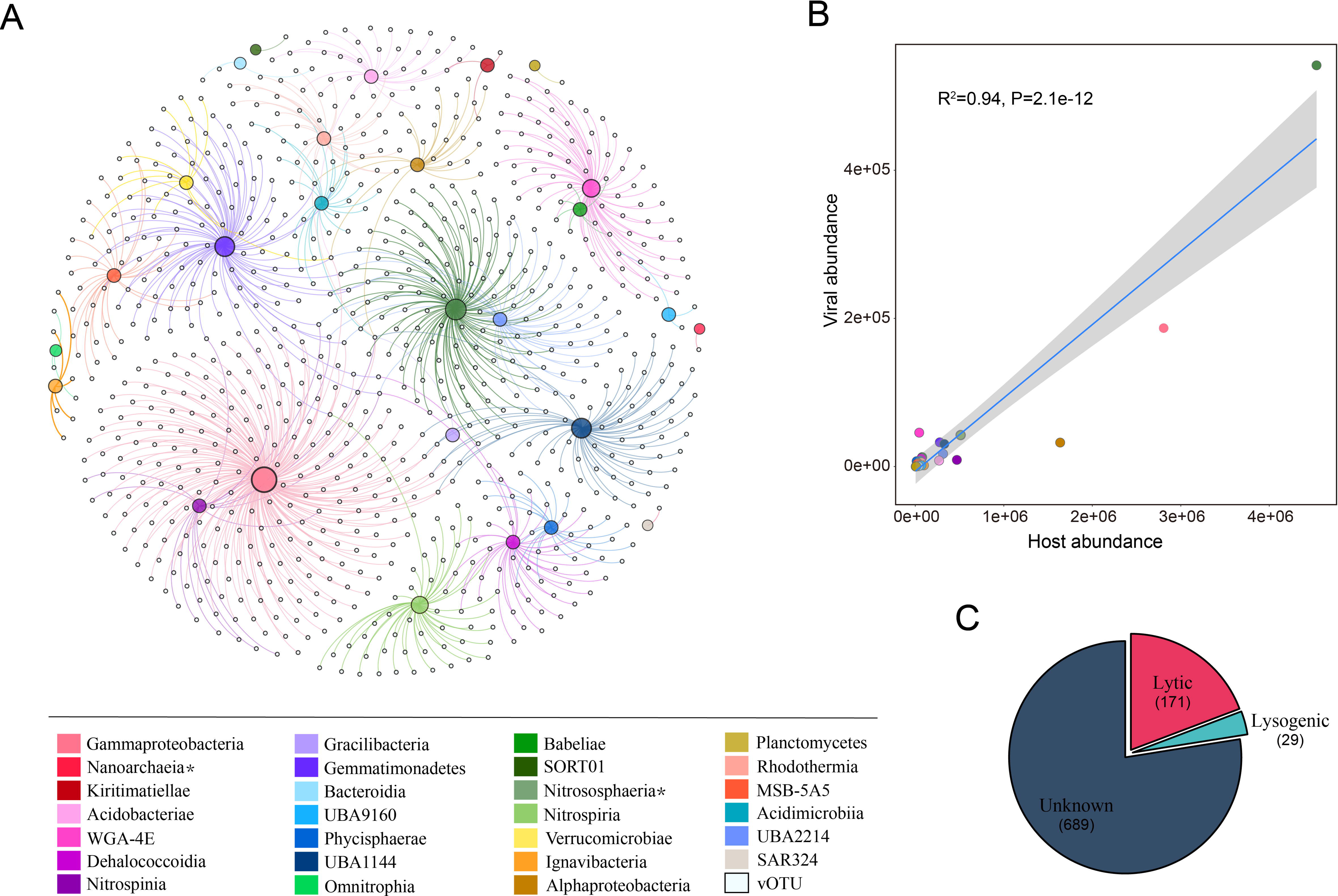
Viruses and their potential host linkage in seamount sediments. **(A)** The connection between viruses and potential hosts is based on the Nucleotide sequence homology, Oligonucleotide frequency (ONF), Transfer RNA (tRNA) match and CRISPR spacer match methods. The size of the host circle depended on the numbers of vOTU. **(B)** Significant Pearson correlation between relative abundances of viruses and their hosts. **(C)** The proportion of host-predicted viral lifestyles. Predicted hosts in **(A)** and **(B)** are indicated in the color bar on the bottom left.

Predicted prokaryotic hosts of vOTUs encompassed two archaeal and twenty-four bacterial classes, representing two archaea and twenty bacterial phyla (Table S6). Of the predicted classes, *Gammaproteobacteria* and *Nitrososphaeria* were the most common hosts, accounting for 25.20% and 11.92% of virus-host linkages, respectively (Table S6). By comparing the data present here to that in the IMG/VR v4 database, two novel viral-bacterial linkages were identified, namely *SORT01* and *Planctomycetes* (Table S6), which implies that these two classes are viral putative hosts, in particular, *SORT01* belongs to *Marinisomatota,* which contains substantial proportions of predicted heme auxotrophs [72], and *Planctomycetes* an important player in global carbon and nitrogen cycles [73]. In addition, some host-virus associations not frequently detected in previous studies were detected in the data presented here, including *Nitrososphaeria*, *Nitrospinia*, *Omnitrophia*, *UBA2214*, *MSB-5A5*, *UBA1144*, *WGA-4E* and *UBA9160* [40, 74, 75]. These results confirm that seamount sediment ecosystems, also contain diverse prokaryotes, with a species complexity similar to that of shallower seamounts [75]. It also implies that seamount sediments are not only diverse in prokaryotes but also have viruses actively infecting these microbes.

### Potential viral auxiliary metabolic genes (vAMGs) associated with biogeochemical cycling and cold shock of host cells

Benthic microorganisms on seamounts occupy diverse habitats and experience a range of environmental conditions [75, 76]. Deep seamounts are usually below the zone of influence of the deep-scattering layer and the biogeography at such depth tends to be less well-characterized. Several studies have highlighted the important contribution of viruses to the carbon, nitrogen, sulfur and phosphorus cycles in sediments [6, 8, 40].

The ORFs identified the Clusters of Orthologous Genes (COGs) function classification through the eggNOG pipeline. Based on the COG classification, 75,057 ORFs (∼78.90% of total ORFs) were clustered into 25 COG categories (Fig. S6). The highest relative abundance of orthologs covered the categories of information storage and processing (LK), metabolism (CE), and cellular processes and signaling (O). Numerous ORFs categorized as having an unknown function (S) were detected at several stations, which implies that seamount sediments still have a large exploration potential. In addition, according to the CAZyme database, the majority of these genes were associated with GlycosylTransferase (GTs) and Glycoside Hydrolases (GHs), which help assemble or break down complex polysaccharides (Fig. S7) [6, 54].

Within the seamount sediment viromes, sixteen different categories of vAMGs involved in the carbon, sulfur, heme, and cobalamin cycle and two vAMGs associated with metal were detected based on the KEGG database (Table S7). For the carbon metabolism genes, only one vAMGs, namely *folD* which was involved in the conversion of the 5,10 Methenyl-THF to 5,10-Methylene-THF, was detected at five out of the twelve sites (Fig. 4A). The Wood–Ljungdahl (WL) pathway is the only carbon fixation pathway in which vAMGs are involved and many anaerobic prokaryotes use this pathway to convert one-carbon substrates (i.e. carbon dioxide or formate) into acetate (Fig. 5B) [77–79].

**Fig. 4.**
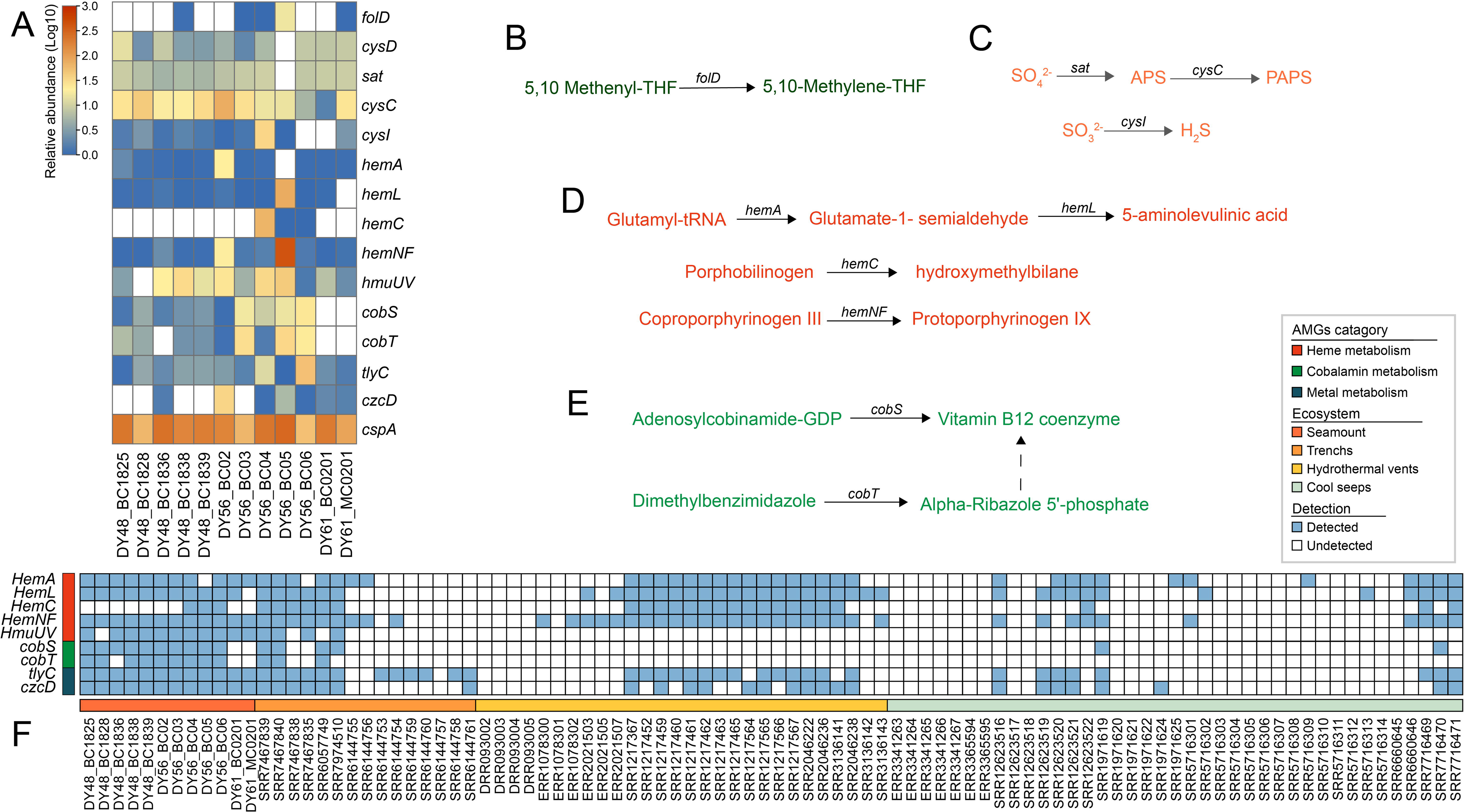
Viral auxiliary aumetabolic genes within the seamount viral genome (SMVG) dataset. **(A)** Heatmap of the relative abundance for different metabolic genes, and results were log10 transformed for description. The cycling pathway of carbon **(B)** sulfur **(C)**, heme biosynthesis **(D)**, and cobalamin biosynthesis (right) **(E)** in seamount sediments. APS: Adenylyl sulfate; PAPS: 3’-Phosphoadenylyl sulfate. **(F)** is the presence or absence of heme, cobalamin and metal-associated vAMGs in four environments. Genes that were not detected in a given sample remained white.

**Fig. 5.**
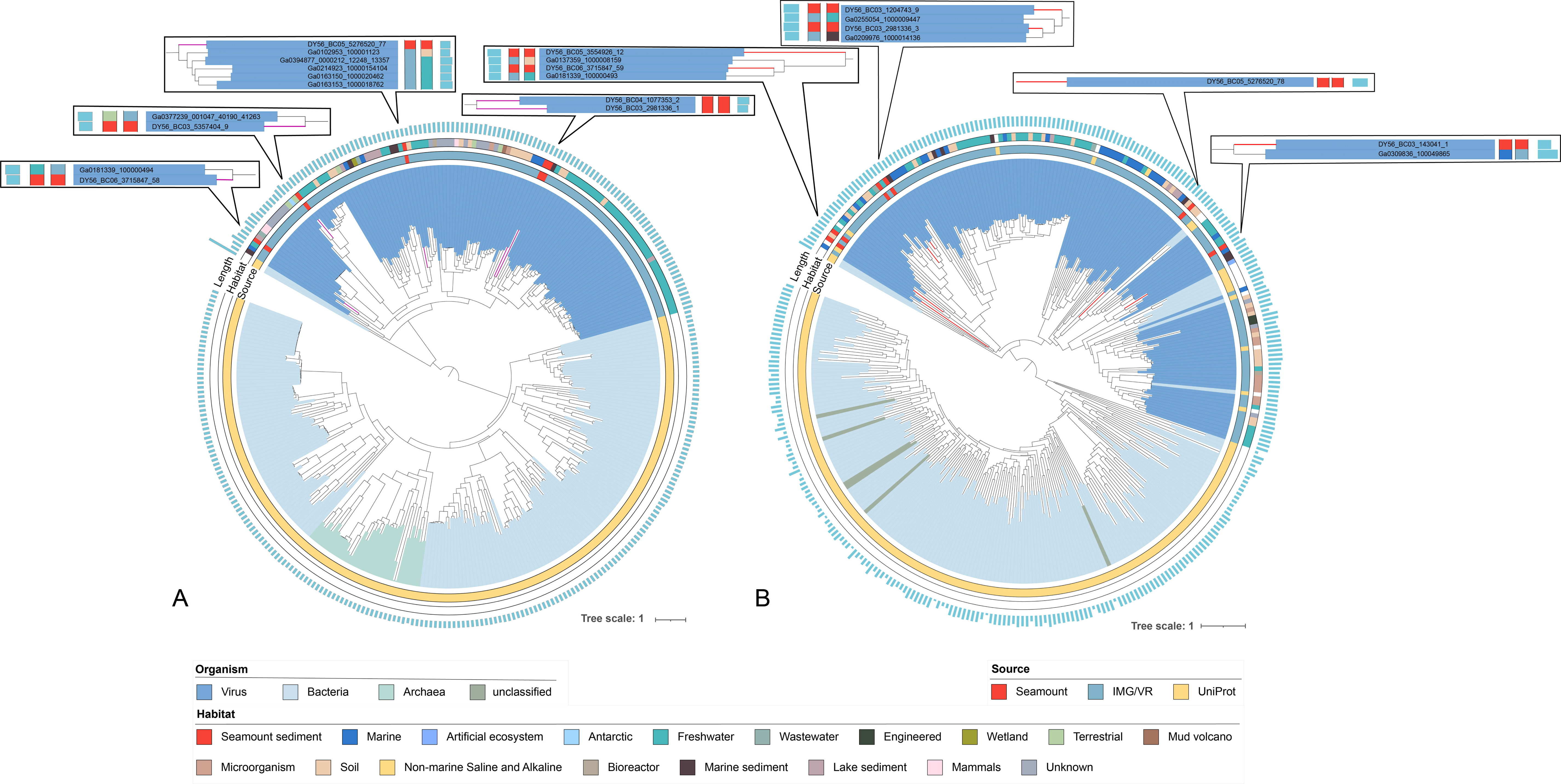
The maximum-likelihood phylogenetic tree of cobaltochelatase *cobS* **(A)** and cobaltochelatase *cobT* **(B)**. *cobS* and *cobT* from seamount sediments are performed by different colors (*cobS*: purple, *cobT*: red). The sequence source are indicated by the inner circle and the habitat of sequences are indicated by the outer circle. The length of each *cobS* sequence ranges from 199 to 538 bp and **t**he length of each *cobT* sequence ranges from 91 to 929 bp.

Pathways for vAMGs that encode sulfur metabolism-related genes were also detected (Fig. 4A). It is noteworthy that most sites contained the function for the assimilatory sulfate reduction pathway (ASR) (Fig. 4C), reflecting that in seamount sediments, viruses tend to complement the hosts’ metabolism through the ASR pathway to accomplish the reduction of sulfate. Furthermore, it was found that *UBA1144* and *Nitrospiria*, typical sulfate-reducing bacteria [80], were associated with 12.26% of the viral contigs, suggesting that the hosts could acquire the corresponding vAMGs which would avoid the energetic costs of the ASR pathway, and the sulfur cycles in seamount sediments are primarily driven by ASR (Fig. 6).

**Fig. 6.**
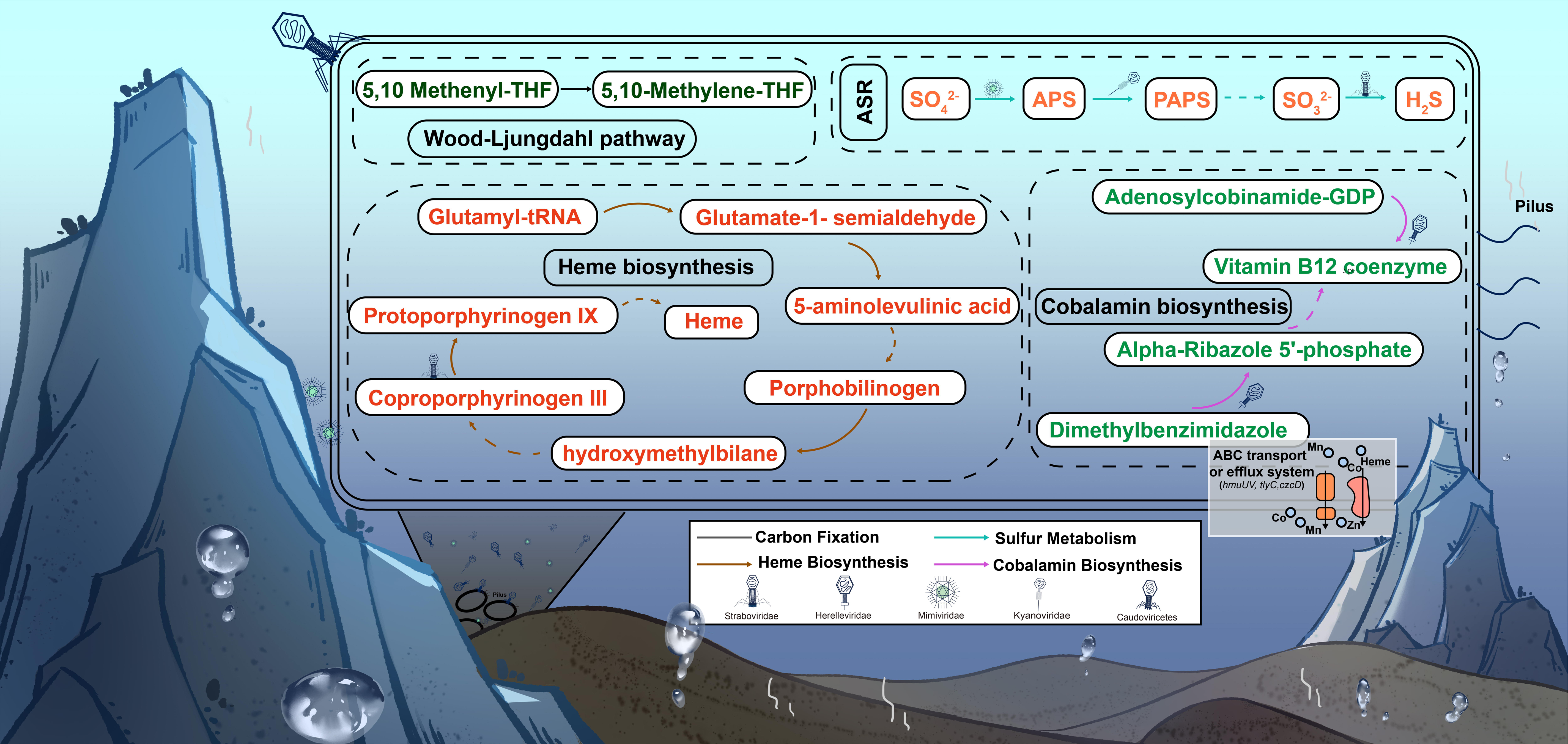
Proposed elemental cycling in seamount sediments with involvement of viruses encoding vAMGs. The figure highlights the proposed elemental cycles of seamount sediments in which vAMGs are involved. Elements including Carbon, Sulfur, Heme, Cobalamin and Metal, have different pathways indicated by different arrow colors. ABC and efflux system (orange and pink) transporters were involved. ASR, assimilatory sulfate reduction.

An AMG encoding a cold shock protein (*cspA*) was also identified at all stations (Table S8). The *cspA* gene functions as an RNA chaperone by destabilizing secondary structures in mRNA, thereby maintaining the single-stranded state of mRNA and concentrating it on transcription and translation [81–83]. The presence of this gene in these samples is consistent with previous studies [84], indicating that viruses might help the host cell to live in the cold deep oceans.

### Comparisons between the seamount sediments and other habitats suggest seamount sediments are a hotspot of AMGs against heme, cobalt and metal metabolisms

Because deep-sea environments are characterized by high hydrostatic pressure, hypoxia and anoxia, extremely high/low temperatures, absence of photosynthesis and nutrient limitation, it is imperative for deep-sea organisms to find adaptions for their survival [85, 86]. Interestingly, two phyla lacking heme biosynthesis were detected in the samples, including *Dependentiae* (class *Babeliae*) and *Patescibacteria* (class *Gracilibacteria*) [72].

Heme (iron-protoporphyrin IX) is an iron-containing tetrapyrrole that is essential for many biochemical reactions, including, oxygenation reactions, oxygen binding and transport, oxidative stress responses, electron transport, oxygen metabolism and xenobiotic detoxification [87–89], heme is also used as a cofactor of hemoproteins. Some prokaryotes lack a heme biosynthesis pathway or contain substantial proportions of predicted heme auxotrophs (e.g., *Dependentiae, Patescibacteria* and *Marinisomatota* in the SMVG data set), these thus require exogenous heme for their growth [72].

Five vAMGs related to heme were detected. While most vAMGs for heme biosynthesis were detected in all twelve samples, the presence of *HemC* was limited to a few specific sites (DY56_BC04, DY56_BC05 and DY56_BC06), indicating that viral-mediated conversion of Porphobilinogen to hydroxymethylbilane might only occur at these three sites. Generally, the formation of 5-aminolevulinic acid (ALA) is considered to be the rate-limiting step of heme biosynthesis, and *HemA* and *HemL* are associated with ALA synthesis [80, 90]. These two genes encoding ALA were abundant at most sites, suggesting that vAMGs can effectively assist in the synthesis of heme in host cells that lack the heme synthesis pathway. Although heme can efficiently help transport iron, heme toxicity is produced when the bacteria’s heme tolerance concentration is exceeded [88, 91]. Heme transport systems-related vAMGs (*HmuUV*) was searched in seamount sediment samples (Fig. 4A). The heatmap shows that the samples had a high relative abundance of *HmuUV*, the heme transport system for ATP-binding protein that powers substrate transports [92]. These results indicate that viruses could provide an additional source, e.g. vAMGs, to supplement the heme biosynthetic pathway.

Intriguingly, two vAMGs encoding cobaltochelatase subunit (*cobS* and *cobT*) were identified in primarily seamount sediments (Fig. 4A). Cobalamin (Vitamin B12), also known as “nature’s most beautiful cofactor”, is used as a cofactor in multiple biological processes, including amino acid metabolism and gene expressions, and is also required by many prokaryotes [93–96]. *cobS* and *cobT* were the subunits that constituted the cobaltochelatase with *cobN*. The three-subunit composition helped cobalt to be inserted into hydrogenobyrinic acid a,c-diamide (HBAD) and form the cobalamin [97, 98]. Therefore, *cobS* and *cobT* genes were analyzed at the amino acid level using IQ-TREE, respectively. Detailed reference sequences including download information and protein names can be found in Table S7 and Table S8. The two phylogenetic trees exhibit the same pattern tree. The *cobS* protein sequences of viruses appear to be evolutionarily distinct from the *cobS* protein sequences of microbes (Fig. 5A), even though some *cobT* protein sequences of viral associated with bacteria, the majority of viral sequences were in the same monophyletic clade, similar to the *cobS* phylogenetic tree (Fig. 5B). However, as all seamount sediment protein sequences cluster with IMG/VR sequences (viral protein sequences), it is concluded that while viral *cobS* and *cobT* originated from microbes and subsequently there was divergent evolution between viruses and microbes. The trend of virus-microbe co-evolution can be inferred in the phylogenic tree of *cobT* with some bacteria sequences inserted in the viral clade, which indicates that horizontal gene transfer (HGT) may have occurred during the evolution of *cobT*. Additionally, the distribution patterns of both vAMGs showed that cobalamin-associated genes have more availability in seamount sediment than in trenches, cool seeps, and hydrothermal vents environments. Furthermore, due to the chromium-rich nature of the sampling site, it is not surprising that the vAMGs were able to assist the host in synthesizing cobalamin. It can thus be speculated that heme and cobalamin play crucial roles as nutrient carriers within deep-sea sediment environments. In conclusion, viruses likely contribute to the biosynthesis of cobalamin in their hosts, since microorganisms are the only source of cobalamin in the ocean and are an essential vitamin for supporting productivity and interactions in the microbial community [94, 99].

Two vAMGs related to the utilization and transport of heavy metals were detected in the SMVG dataset, including, magnesium and cobalt exporter (*tlyC*) and cobalt-zinc-cadmium efflux system protein (*czcD*), which related to the cobalt metabolism (Fig. 4A). Then ten noteworthy vAMGs found in trenches, cool seeps, and hydrothermal vents environments were underrepresented (Fig. 5). Although vAMGs associated with heme and metal metabolism were detected more frequently in the other three environments, primarily in sediments of trenches, plumes of hydrothermal vents and sediment of cool seeps, when compared to cobalamin-associated vAMGs (Supplement Table S8), there were significant differences between seamount and most other sites and this suggests that the above metabolisms were more important biogeochemical processes in seamount sediments than in other environments.

The vAMGs observed in the seamount sediments indicate that viruses in the deep ocean not only contribute to prokaryotic metabolism and nutrient cycling but also assist the host cells in adapting to nutrient-limiting and cold habitats (Fig. 6).

## Conclusions

Due to the challenges associated with sampling deep-sea sediments and the difficulties in laboratory cultivation of microbial communities and their corresponding viruses, the genome, diversity, evolution and ecological roles of viruses remain largely unexplored in deep-sea marine sediments associated with seamounts. Here, for the first time, the diverse and unique viral assemblages inhabiting the seamount sediments are identified, most of which are different from other cultured and environmental viral communities. Novel virus-host linkages were identified from the SMVG dataset, e.g. *SORT01* and *Planctomycetes*. Viruses encoding vAMGs, encompassing genes related to the metabolism of carbon, sulfur, heme, cobalamin and heavy metals, suggest potentially important roles of viruses in the biogeochemical and heavy-metals of seamount sediments in the Northwest Pacific Ocean (Fig. 6). This study sheds light on the diversity and ecological significance of benthic viruses, and provides a baseline to incorporate viruses in model of deep-sea seamount sediment ecosystems. With the continuous improvement of sampling and *in-situ* cultivation technologies, and increasing numbers of scientific cruises focusing on deep-sea seamounts, a more comprehensive understanding of viruses in the deep-sea seamounts is expected in the near future.

## Data availability

The raw sequence data reported in this paper have been deposited in the Genome Sequence Archive (Genomics, Proteomics & Bioinformatics 2021) in National Genomics Data Center (Nucleic Acids Res 2022), China National Center for Bioinformation / Beijing Institute of Genomics, Chinese Academy of Sciences (GSA: CRA012520) that are publicly accessible at https://ngdc.cncb.ac.cn/gsa/s/89C84cZv.

## Acknowledgments

Grant support was provided by the high-performance server at the Center for High Performance Computing and System Simulation, Pilot National Laboratory for Marine Science and Technology (Qingdao, China), a high-performance computing cluster operated by the Institute of Evolution and Marine Biodiversity, and the high-performance servers of Marine Big Data Center of Institute for Advanced Ocean Study of Ocean University of China. We thank the captain and crew of the cruises DY48, DY56 and DY61, for their assistance with the shipboard experiment.

## Authors’ contributions

Q.L., M.W., and Y.T.L. conceived and designed the experiment. Y.C., and C.G. methodology, formal analysis, writing, and original draft preparation. M.Y.L., and Y.C. DNA extraction and metagenomic dataset, Y.C. F.Y.G., and H.Y. participated in statistical analysis. Q.L., A.M., M.W., and Y.T.L. critically revised the manuscript. All authors revised and commented on the article and approved the final version.

## Ethics approval and consent to participate

Not applicable

