## Supplemental information for "Diversity and ecological potentials of viral assemblages from the seamount sediments of the Northwest Pacific Ocean"

**Supplemental Methods:**

**DNA extraction and sequencing**

To the best of our knowledge, this is the first study to describe the viral communities buried in seamount sediments of the western Pacific Ocean using metagenomic methods However, because of insufficient biomass of the collected samples, MDA amplification, a method for DNA amplification, was applied in this study [1]. MDA has a performance bias in preferentially amplifying genomes of ssDNA, extreme GC, and uneven coverage [2]. Therefore, these data were processed with sensitivity and an attempt was made to minimize biased descriptions in interpreting the results. However, it needs to be emphasized that all identified species were present in our samples, the method used only influenced the relative abundance and not the presence/absence of a taxon. In addition, several features suggested a further bias in the MDA in this study, including: (1) the recognized best choice of MDAkits was used to minimize the bias of background contaminating DNA, namely the QIAGEN REPLI-g Mini Kit [3]; (2) Regarding past studies, the effects of MDA amplification induced has been considered random, and the other unknown factors must be participating [4]; (3) In addition, although ssDNA viruses were found in the trench, their relative abundance was low, which means that MDA method might exhibit a different pattern in environmental contigs [5]. The MDA method has been considered a better choice in metagenomic surveys, especially when the volume of samples is insufficient. In the future, the library preparation based on the low-input library preparation without MDA for the high-throughput sequencing (such as NexteraXT and NebNext UltraII kits) may be able to provide better results.

**Supplemental Figures:**

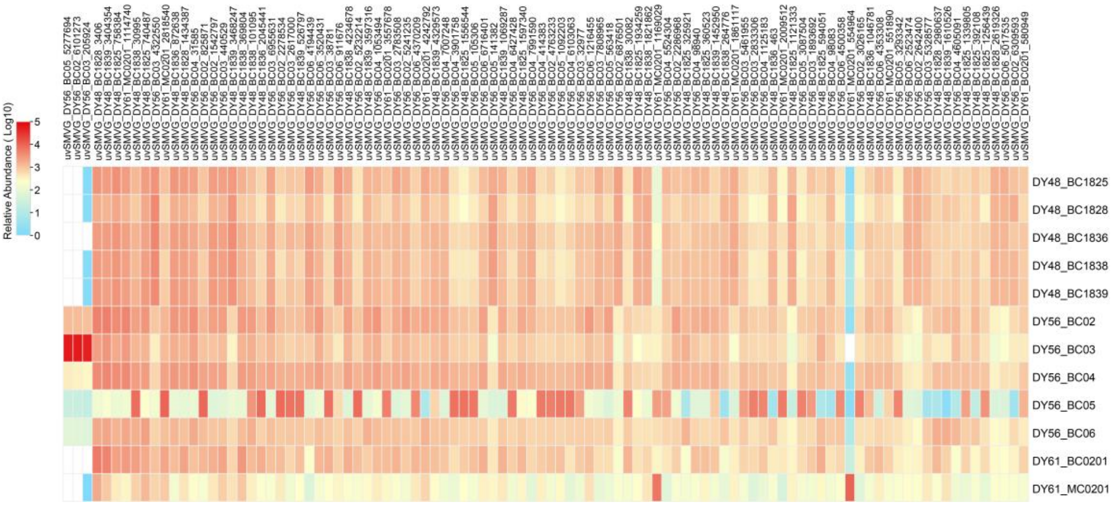


**Fig. S1.** The most 100 abundant vOTUs in the seamount sediments, and results were log10 transformed for description.

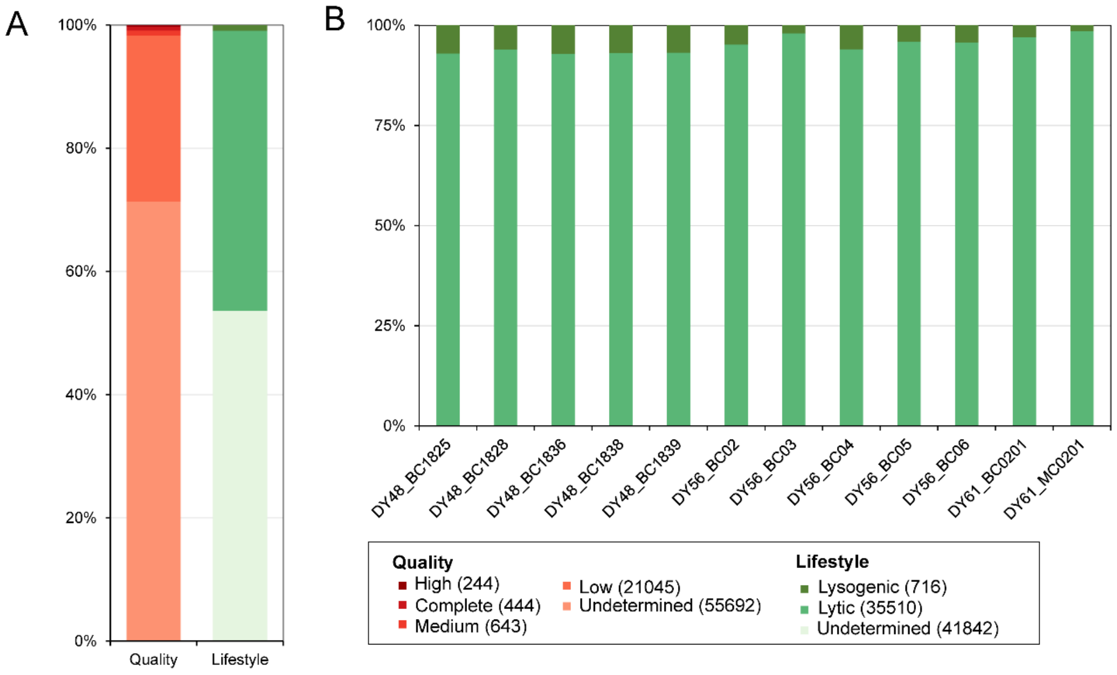


**Fig. S2.** Overview of viral communities in seamount sediments. **(A)** Bar charts showing the quality and lifestyle of SMV viral contigs. **(B)** Bar charts showing the number of lytic and lysogenic viral contigs in each station.

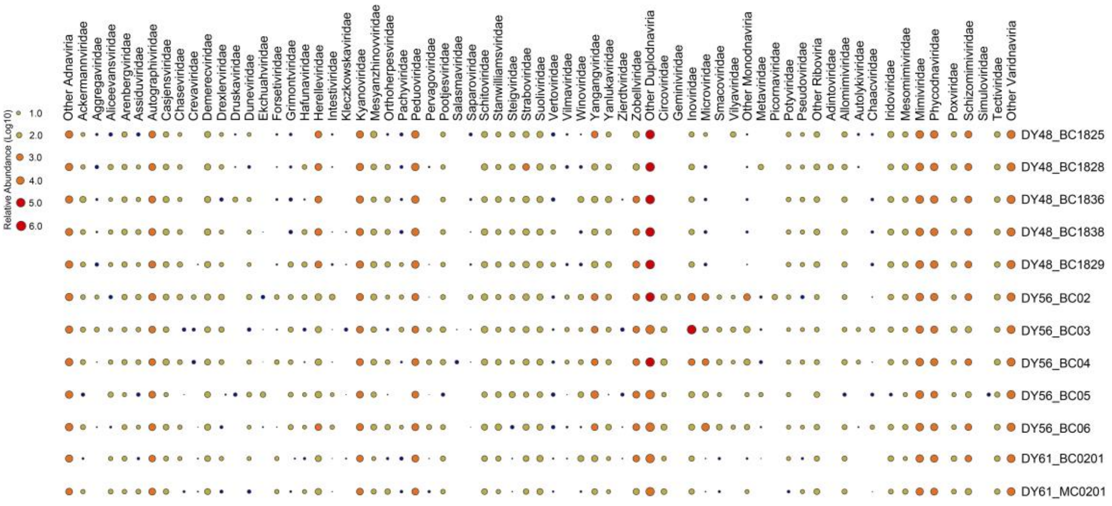


**Fig. S3.** Heatmap of the all viral taxonomical assignment (family level). The relative abundance was calculated with TPM, and the results were log10 transformed for description.

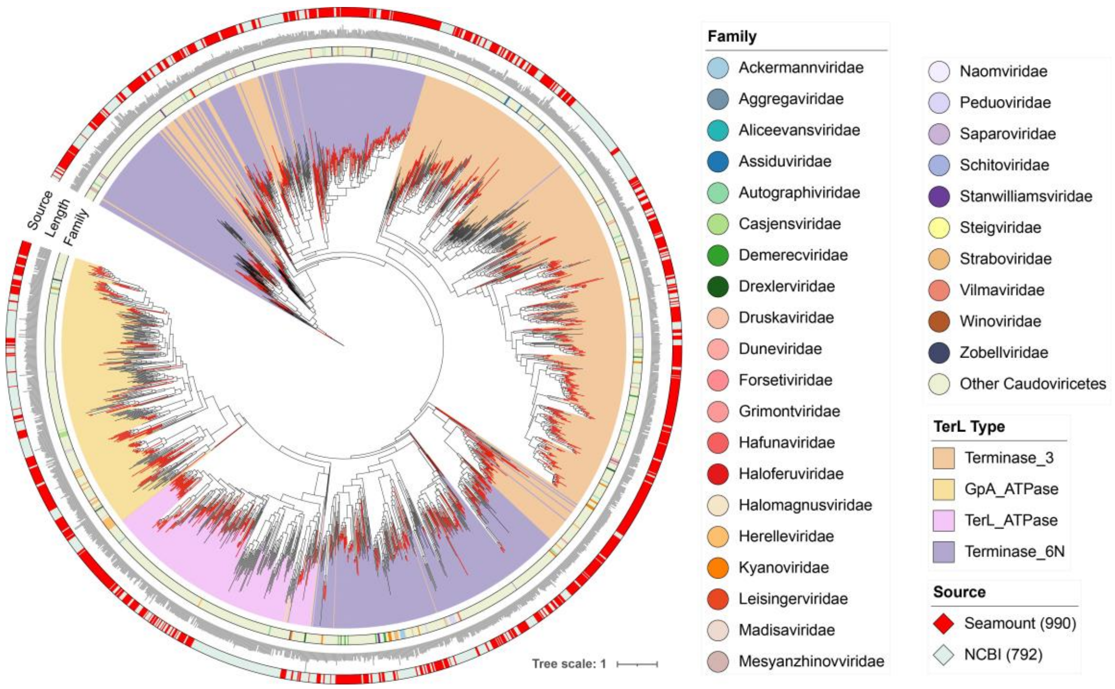


**Fig. S4.** The maximum-likelihood phylogenetic tree of terminase large subunits (TerL) of *Caudoviricetes*. Different TerL domains are performed by different colors. The family levels were indicated by the inner circle and the source viral TerL domain sequences were indicated by the outer circle. The length of each TerL domain sequence ranges from 81 to 1284 bp.

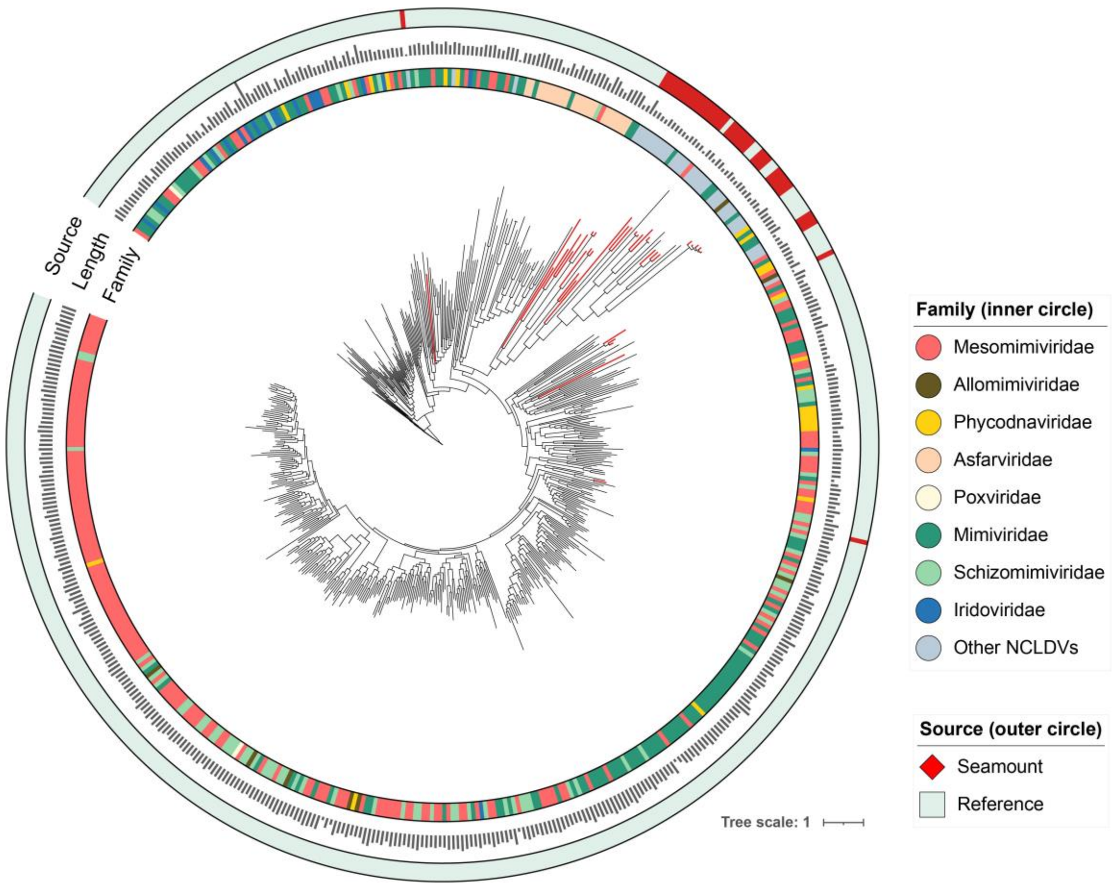


**Fig. S5.** The maximum-likelihood phylogenetic tree of *polB* of NCLDVs. The family levels were indicated by the inner circle and the source viral *polB* sequences were indicated by the outer circle. The length of each *polB* domain sequence ranges from 177 to 3158 bp.

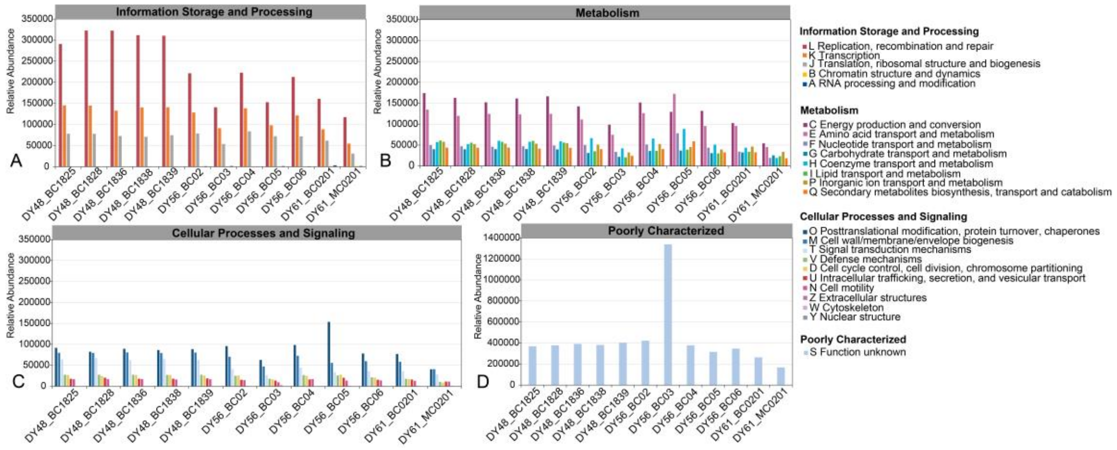


**Fig. S6.** The relative abundance of the NCBI COG functional categories of genes annotated by eggNOG-Mapper. The relative abundance is calculated with TPM.

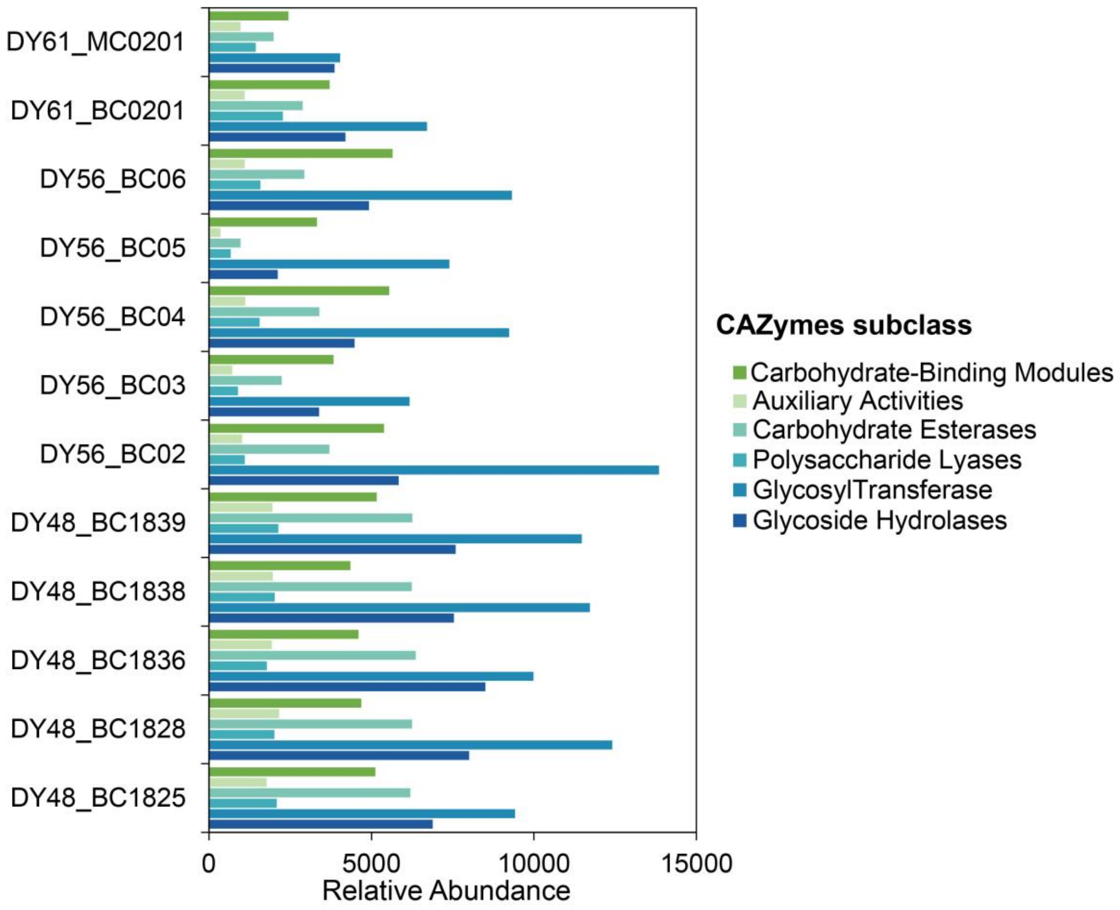


**Fig. S7.** The relative abundance of virus-encoded CAZyme genes in the data set. The relative abundance is calculated with TPM.
